# miR-200c suppresses stemness and increases cellular sensitivity to trastuzumab in HER2+ breast cancer

**DOI:** 10.1101/130682

**Authors:** Cailu Song, Jin Wang, Hua Wang, Peng Liu, Longzhong Liu, Lu Yang, Hailin Tang, Xiaoming Xie

## Abstract

Resistance to trastuzumab remains a major obstacle in HER2-overexpressing breast cancer treatment. miR-200c is important for many functions in cancer stem cells (CSCs), including tumor recurrence, metastasis and resistance. We hypothesized that miR-200c contributes to trastuzumab resistance and stemness maintenance in HER2-overexpressing breast cancer. In this study, we used HER2-positive SKBR3, HER2-negative MCF-7, and their CD44^+^CD24^-^ phenotype mammospheres SKBR3-S and MCF-7-S to verify. Our results demonstrated that miR-200c was weakly expressed in breast cancer cell lines and cell line stem cells. Overexpression of miR-200c resulted in a significant reduction in the number of tumor spheres formed and the population of CD44^+^CD24^-^ phenotype mammospheres in SKBR3-S. Combining miR-200c with trastuzumab can significantly reduce proliferation and increase apoptosis of SKBR3 and SKBR3-S. Overexpression of miR-200c also eliminated its downstream target genes. These genes were highly expressed and positively related in breast cancer patients. Overexpression of miR-200c also improved the malignant progression of SKBR3-S and SKBR3 in vivo. miR-200c plays an important role in the maintenance of the CSC-like phenotype and increases drug sensitivity to trastuzumab in HER2+ cells and stem cells.

**Summary statement:** miRNAs are critical in stemness maintenance and drug resistance. These data link maintenance of the stemness-related phenotype and the sensitivity of HER2+ breast cancer to miR-200c in response to trastuzumab.

## Introduction

HER2 gene amplification occurs in 20-25‥ of breast cancers and is associated with high relapse and poor prognosis rates (Bose et al., 2013; Slamon et al., 1987; Yarden and Pines, 2012). Trastuzumab (Herceptin) is a monoclonal antibody that inhibits downstream signaling of intracellular transduction by targeting the extracellular domain of the HER2 gene. Although trastuzumab is effective, the efficiency is only approximately 26%, even in HER2-overexpressing breast cancer patients. The median remission is approximately 9 months, and the majority of patients acquire resistance to trastuzumab within 1 year (Rexer et al., 2013; Walsh et al., 2013). In combination with another HER2-targeted drug Lapatinib, a tyrosinase inhibitor, the treatment remains ineffective in approximately half of patients. Trastuzumab therapy may increase the risk of brain metastases in some patients with breast cancer (Cortes et al., 2015; Del Mastro et al., 2012). Trastuzumab treatment costs approximately US$45,000 a year in China, which is an enormous financial burden to families of breast cancer patients (Shiroiwa et al., 2008). To overcome resistance and improve the efficacy of trastuzumab treatment in HER2-overexpressing breast cancer patients, we must perform a thorough inquiry into the mechanisms of trastuzumab resistance and develop new effective treatment programs.

A variety of potential molecular mechanisms of resistance to trastuzumab have previously been published (Berns et al., 2007; Nagata et al., 2004; Reim et al., 2009; Zhang et al., 2011), and the vast majority involve the biological functions of breast cancer stem cells (BCSCs) (Castagnoli et al., 2016; Reim et al., 2009). Cancer stem cells (CSCs) are a rare fraction of tumor cells that have the abilities of self-renewal, unlimited proliferation and multi-potent differentiation (Dalerba and Clarke, 2007). Like normal tissues, tumor tissues are composed of a variety of heterogeneous tumor cells and originate from corresponding stem cells. CSCs have been isolated from a variety of tumors, such as leukemia, lung cancer, and breast cancer (Dietrich et al., 2014; Ho et al., 2007; Marotta et al., 2011). There are two recognized methods of separating BCSCs from breast cancer patients and cell lines: one is surface phenotypic marker screening, and the other is Hoechst 33342 dye exclusion. CD44^+^CD24^-^ is a well-known surface maker for the isolation and identification of BCSCs in breast cancer tissues and cell lines (Al-Hajj et al., 2003; Marotta et al., 2011).

There is mounting evidence that CSCs are responsible for tumor formation, infinite growth, recurrence and metastasis. CSCs have congenital resistance features. Conventional drug therapy, including chemotherapy drugs, radiotherapy drugs and targeted therapy drugs, can kill only the active non-stem cells, whereas residual CSCs can eventually lead to tumor recurrence and metastasis. CSCs are the root cause of drug resistance and treatment failure (Hong et al., 2009; Lemoli et al., 2009). The mechanisms of drug resistance in CSCs include overexpression of ATP-binding cassette transporters, over-activation of cell detoxification enzymes, abnormal activation of cell survival and apoptosis-related signal transduction pathways, the protective effect of tumor niches on tumor stem cells, and that most CSCs are in a quiescent phase (Bardelli and Siena, 2010; Gottesman, 2002; Lin et al., 2010). By intervening in these processes, we may reverse resistance to trastuzumab and improve the survival and prognosis of breast cancer patients.

MicroRNAs (miRNAs) are a class of endogenous small RNAs that regulate many crucial biological processes in cancer pathogenesis and progression. miRNAs are highly conserved and specific. By binding to the 3’-untranslated region (UTR) of target messenger RNA (mRNAs), miRNAs can regulate gene expression by inhibiting translation and inducing degradation of mRNAs (Calin and Croce, 2006). miRNAs act as either oncogenes or tumor suppressors in cancer management (Bartel, 2004). Thus far, many aberrantly expressed miRNAs have been discovered in different cancers. The miR-200 family is one family of these miRNAs. The miR-200 family consists of miR-200a, miR-200b, miR-200c, miR-141 and miR-429. According to their location on two different chromosomes, the miR-200 family can be divided into two genetically different subfamilies that have essentially the same seed sequence, namely the miR-200c/miR-141 and miR-200a/miR-200b/miR-429 gene clusters (Bracken et al., 2014).

As a tumor suppressor, miR-200c has caused extensive concern. Several studies associated miR-200c and its target mRNAs with the establishment, maintenance and regulation of the CSC phenotype. Down-regulation of miR-200c is relevant for stem cell functions in cancer, including self-renewal, clonal expansion, and differentiation (Shimono et al., 2009). miR-200c inhibits tumor growth, differentiation and self-replication of CSCs by targeting TUBB3 and restoring sensitivity to microtubule-targeting drugs (Liu et al., 2013). Members of the miR-200 family control the EMT process and sensitivity to EGFR therapy in bladder cancer (Adam et al., 2009). Down-regulation of miR-200c expression is a marker of tumor invasion and drug resistance in female genital tumors, such as ovarian cancer, cervical cancer and breast cancer (Cochrane et al., 2010). Therefore, exploring the role of miR-200c in BCSCs may help us understand the mechanism of drug resistance.

Increasing evidence suggests that the Notch, Wnt and Hedgehog signal transduction pathways are the most important pathways in stemness maintenance and drug resistance of CSCs (Takebe et al., 2011; Zardawi et al., 2009). The Notch family consists of transmembrane signal proteins, and their main function is to determine the direction of cell development. Notch is highly expressed in tissue stem cells and precursor cells. Notch promotes stem-like mammosphere formation and increases the recurrence of dormant tumor cells following HER2/neu-targeted therapy of BCSCs (Abravanel et al., 2015; Grudzien et al., 2010). Abnormal activation of the Wnt pathway interferes with the survival, self-renewal, proliferation and drug insensitivity of BCSCs, causes an increase in the proportion of stem cells, induces a conversion from mammary stem cells into BCSCs, and ultimately results in the recurrence and malignant progression of breast cancer (Ramachandran et al., 2014; Xu et al., 2016). The Hedgehog signaling pathway is highly activated in BCSCs, and inhibition of the Hedgehog pathway significantly inhibits the proliferation of CSCs and enhances their sensitivity to drugs (Tanaka et al., 2009; Yoo et al., 2011).

In addition, Jagged1, ZEB1 and Bmi1 are highly critical genes in these three signaling pathways. They are the direct target genes of the miR-200 family in many cancers, including breast cancer (Korpal et al., 2008; Liu et al., 2014; Vallejo et al., 2011). Based on the above, BCSCs are the key factor responsible for trastuzumab resistance in HER2-overexpressing breast cancer. miR-200c may suppress stemness and increase the sensitivity to trastuzumab in HER2+ breast cancer cells and stem cells. The study of the mechanism may be helpful for targeting BCSCs that are not successfully killed by trastuzumab and may improve the survival and prognosis of HER2-overexpressing breast cancer patients.

## Materials and Methods

### Clinical samples

All human breast cancer tumor samples were obtained from randomly selected cancer patients at the Sun Yat-Sen University Cancer Center (SYSUCC), Guangzhou, China. The diagnoses of breast cancer were pathologically confirmed in all cases. The patients had not received any prior chemotherapy. Tissue samples were collected intraoperatively. Written informed consent was obtained from all eligible patients who participated in the study before or after surgery, and all protocols were reviewed and approved by the Joint Ethics Committee of SYSUCC and used according to the ethical standards as formulated in the Helsinki Declaration.

### Adherent cell and tumor sphere culture

All adherent breast cancer cell lines used in this study were purchased from American Type Culture Collection (Manassas, VA, USA). MCF-7, T47D, BT-474, SKBR3, MDA-MB-453, MDA-MB-468, HCC-38, MDA-MB-231 and BT-549 cells were maintained in DMEM with 10% fetal bovine serum (FBS), penicillin (100 U/mL), and streptomycin (100 μg/mL) (Invitrogen, Carlsbad, CA) and incubated at 5% CO2 at 37°C. MCF-10A cells were cultured in DMEM/F12 supplemented with 5% donor horse serum, epidermal growth factor (EGF, 20 ng/ml), insulin (10 μg/ml), hydrocortisone (0.5 μg/ml), penicillin (100 units/ml) and streptomycin (100 μg/ml) (Invitrogen).

CD44^+^CD24^-^ phenotype cells were detected and sorted using flow-cytometric analysis. CD44^+^CD24^-^ flow-sorted cells were cultured to form spheres in ultralow attachment dishes (BD Biosciences, San Diego, CA) at a density of 1,000 cells per milliliter for 5 days. Serum-free suspension culture stem cell spheres (MCF-7-S, T47D-S, BT-474-S, SKBR3-S, MDA-MB-453-S, MDA-MB-468-S, HCC-38-S, MDA-MB-231-S, BT-549-S and MCF-10A-S) were grown in DMEM/F12 supplemented with 1% B27, EGF (20 ng/ml), basic fibroblast growth factor (bFGF) (10 ng/ml), and insulin (10 mg/ml) (Invitrogen). The spheres were counted and collected 5 days after being planted. They were mechanically dissociated with a 25 G needle (BD) and reseeded in fresh medium.

### Mammosphere formation assay

Cells were plated in 6-well plates for 24 h and transfected. After incubation for 48 h, cells were harvested and plated at 104 cells/well in ultra-low-attachment six-well plates in serum-free DMEM/F12 supplemented with B27 (1:50), EGF (20 ng/ml), bFGF (10 ng/ml) and insulin (10 mg/ml). Mammospheres were counted after 5 days of culture using a Nikon Eclipse TE2000-S microscope (Japan) and photographed with Meta Morph.

### Transient and stable overexpression of miR-200c

Recombinant lentivirus vectors encoding miR-200c (miR-200c) or negative control (Control) (pEZX-MR03, HIV based, GeneCopoeia, USA) were used for transient or stable overexpression of miR-200c. All cells used in this experiment were stably transfected with recombinant lentivirus vectors encoding miR-200c or its control, except the MTT and apoptosis assays (transient transfection). The overexpression plasmid and its control were transfected into cells using Lipofectamine 2000 (Invitrogen) following the manufacturer’s instructions. At 6 h after transfection, the culture medium was changed to DMEM supplemented with 2% FBS or DMEM/F12. At 24 h or 48 h after the transfection, the cells were harvested for the isolation of total RNA or protein. The stable transfected cells were additionally infected in the presence of polybrene (4 mg/ml) and selected with puromycin (1 mg/ml).

### RNA extract and qRT-PCR

Total RNA from cells was extracted with TRIzol reagent (Invitrogen) following the manufacturer’s protocol. Reverse transcription and quantitative real-time PCR analysis (qRT-PCR) reactions were performed using a qSYBR-Green-containing PCR kit (Qiagen, Germantown, USA) and a BioRad IQTM5 Multicolor Real-Time PCR Detection System (USA). The primers for Jagged1, ZEB1 and Bmi1 mRNA for qRT-PCR detection were synthesized by Invitrogen. Each experiment was performed three times in triplicate.

### Protein isolation and western blotting

Protein was extracted from cells lines using RIPA lysis buffer with proteinase inhibitors (Thermo, Rockford, USA). The Protein BCA Assay Kit (Thermo) was used to measure the concentration of protein in the lysates. Each lane was loaded with 20 μg protein mixed with 2× SDS loading buffer (Thermo). The protein was separated by 12% SDS-polyacrylamide gel electrophoresis and then transferred to polyvinylidene difluoride membranes (Millipore, Bedford, USA). The membranes were incubated with 5% skim milk powder at room temperature for 1 h to block nonspecific binding. Subsequently, the membranes were incubated with antiserum containing antibodies against Jagged1, ZEB1 and Bmi1 (CST, Massachusetts, USA), at 4°C for 12 h. To visualize the target proteins, a 1:5000 dilution of peroxidase-conjugated secondary antibody and ECL western blotting detection reagents (GE, USA) were used. A Bio Image Intelligent Quantifier 1-D was used to quantify the proteins (Version 2.2.1, Nihon-BioImage Ltd., Japan). An anti-GAPDH antibody was used as a protein loading control (Boster, Wuhan, China).

### MTT assay

Cell viability analysis was preformed according to the methods using 3-(4, 5-dimethylthiazol)-2, 5-diphenyltetrazolium bromide (Tang et al., 2011). SKBR3 and SKBR3-S cells were divided into mock (negative control), Trastuzumab (10 μg/ml), Control (transiently transfected with lentiviral vector coding control), miR-200c (transiently transfected with lentiviral vector coding miR-200c), and Trastuzumab+miR-200c groups. To assess the incorporation of BrdUrd, all cells in the treated and control groups were incubated with 10 nM BrdUrd in culture media for 16 h, and then the cells were fixed with cold methanol and acetone (1:1).

### Apoptosis rate determined by flow cytometry

SKBR3 and SKBR3-S were divided into five groups as in the MTT assay. Cells at 5×105 cells/ml were inoculated into 6-well culture plates and incubated at 37°C. After culture for 72 h, cells were collected after digestion with 0.25% trypsin, washed with PBS three times, and resuspended in 500 ml binding buffer (10 mM HEPES/NaOH, 140 mM NaCl, 2.5 mM CaCl2, PH 7.4). FITC-labeled Annexin V (50 mg/ ml, 5 ml) and PI (50 mg/ml, 5 ml) were added, followed by incubation at room temperature in the dark for 30 min. The apoptosis rate was immediately measured by flow cytometry.

### Sort BCSCs by flow-cytometry

Serum-free suspension culture stem cell spheres, cells were collected and rinsed twice with PBS. Then cells were stained with phycoerythrin-conjugated anti-human CD24 antibody (Invitrogen) and FITC-conjugated anti-human CD44 antibody (Invitrogen) for 30 min at room temperature according to the manufacturer’s instructions. The labeled cells were then analyzed using a FACSCalibur flow cytometer and Cell Quest software (BD).

### Human breast cancer xenograft assay

CD44^+^CD24^-^ human BCSCs were isolated by flow cytometry. BCSCs (SKBR3-S and MCF-7-S) and breast cancer cells (SKBR3 and MCF-7) were infected with 20 MOI of miR-200c expressing lentivirus or control lentivirus by spin infection for 2 hours followed by incubation at 37 °C for 2 hours. After establishing stable cell lines expressing miR-200c or control, the cells were collected and mixed with Matrigel (BD). Either 5,000 or 10,000 cells were injected into the mammary fat pad of female Balb/c mice. All experiments were performed under the approval of the Administrative Panel on Laboratory Animal Care of SYSUCC.

### Statistical analysis

Each experiment was repeated three times, and the results are presented as the mean ± SE. The statistical analysis was performed using Student’s t test and Pearson correlation analysis. Fisher’s exact test was used to analyze the significance of in vivo experiment results. The significance level was set at p < 0.05. All statistical analyses were performed using SPSS 17.0 software.

## Results

### miR-200c was weakly expressed in breast cancer cell lines and CD44^+^CD24^-^ phenotype cell line stem cells

We collected nine breast cancer cell lines (MCF-7, T47D, BT-474, SKBR3, MDA-MB-453, MDA-MB-468, HCC-38, MDA-MB-231 and BT-549) and one mammary epithelial cell line (MCF-10A). We also collected nine CD44^+^CD24^-^ phenotype breast cancer cell line stem cells (MCF-7-S, T47D-S, BT-474-s, SKBR3-S, MDA-MB-453-S, MDA-MB-468-S, HCC-38-S, MDA-MB-231-S and BT-549-S) and one CD44^+^CD24^-^ phenotype mammary epithelial cell line stem cells (MCF-10A-S), which were serum-free suspension cultured and then sorted by flow cytometry. qRT-PCR was used to detect the expression of miR-200c in these cells. We found that miR-200c was downregulated in both breast cancer cell lines and CD44^+^CD24^-^ phenotype breast cancer cell line stem cells (Figure 1A, 1B).

**Figure 1.**
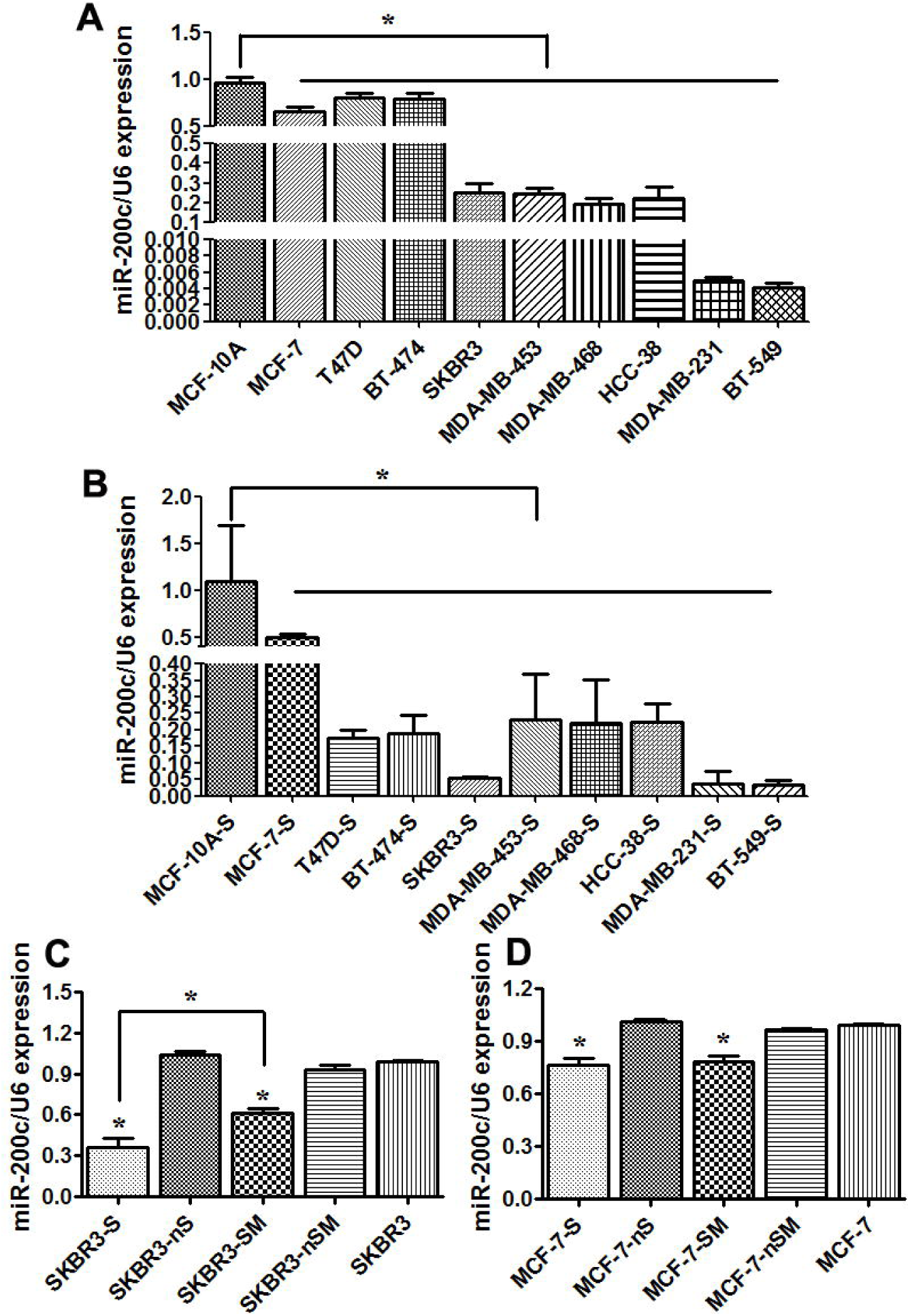
miR-200c was weakly expressed in breast cancer cell lines and CD44^+^CD24^-^ phenotype cell line stem cells. (A) miR-200c expression in breast cancer cell lines and mammary epithelial cells. miR-200c was downregulated in MCF-7, T47D, BT-474, SKBR3, MDA-MB-453, MDA-MB-468, HCC-38, MDA-MB-231 and BT-549 cells (*p<0.05 vs. MCF-10A). (B) miR-200c expression in breast cancer cell line stem cells and mammary epithelial stem cells. miR-200c was downregulated in CD44^+^CD24^-^ phenotype MCF-7-S, T47D-S, BT-474-s, SKBR3-S, MDA-MB-453-S, MDA-MB-468-S, HCC-38-S, MDA-MB-231-S and BT-549-S cells, which were serum-free suspension cultured and then sorted by flow cytometry (*p<0.05 vs. MCF-10A-S). (C) miR-200c expression in CD44^+^CD24^-^ phenotype BCSCs and the remaining non-CD44^+^CD24^-^ phenotype breast cancer cells from SKBR3 that were sorted by flow cytometry (SKBR3-S/SKBR3-nS), suspension microspheres and the remaining cells after filtration of the microspheres from SKBR3 that only serum-free suspension cultured and not sorted by flow cytometry (SKBR3-SM/SKBR3-nSM) and normal adherent cells of SKBR3. miR-200c was downregulated in SKBR3-S and SKBR3-SM (*p<0.05 vs. SKBR3). The expression of miR-200c was much lower in CD44^+^CD24^-^ phenotype SKBR3-S than suspension microsphere SKBR3-SM (*p<0.05 vs. SKBR3-SM). (D) miR-200c expression in HER2-CD44^+^CD24^-^ phenotype BCSCs and theremaining non-CD44^+^CD24^-^ phenotype breast cancer cells from MCF-7 that were sorted by flow cytometry (MCF-7-S/MCF-7-nS), suspension microspheres and the remaining cells after filtration of the microspheres from MCF-7 that only serum-free suspension cultured and not sorted by flow cytometry (MCF-7-SM/MCF-7-nSM) and normal adherent cells of MCF-7. miR-200c was downregulated in MCF-7-S and MCF-7-SM (*p<0.05 vs. MCF-7). There were no significant differences between the expression of miR-200c in MCF-7-S and MCF-7-SM (*p>0.05 vs. MCF-7-SM).

Then, we selected HER2+ SKBR3 and HER2- MCF-7 cells for further study. qRT-PCR was used to detect the expression of miR-200c in CD44^+^CD24^-^ phenotype BCSCs and the remaining non-CD44^+^CD24^-^ phenotype breast cancer cells that were sorted by flow cytometry (SKBR3-S/SKBR3-nS, MCF-7-S/MCF-7-nS), suspension microspheres and the remaining cells after filtration of the microspheres that only serum-free suspension cultured and not sorted by flow cytometry (SKBR3-SM/SKBR3-nSM, MCF-7-SM/MCF-7-nSM), normal adherent cells of SKBR3 and MCF-7. By comparing with SKBR3 and MCF-7 cells, we found that miR-200c was downregulated in SKBR3-S, SKBR3-SM and MCF-7-S, MCF-7-SM (Figure 1C, 1D). The expression of miR-200c was much lower in CD44^+^CD24^-^ phenotype SKBR3-S than in the suspension microspheres SKBR3-SM (Figure 1C). However, there were no significant differences between the expression of miR-200c in MCF-7-S and MCF-7-SM (Figure 1D). **Overexpression of miR-200c resulted in a decrease in the number of tumor spheres and proportion of CD44^+^CD24^-^ phenotype cells in HER2+ BCSCs**

To further confirm the effect of miR-200c on tumor sphere formation in HER2+ breast cancer cells, CD44^+^CD24^-^ phenotype SKBR3-S and MCF-7-S cells were stably transfected with lentiviral vector encoding miR-200c and its negative control. The transfection of miR-200c was successful (Figure 2A).

**Figure 2.**
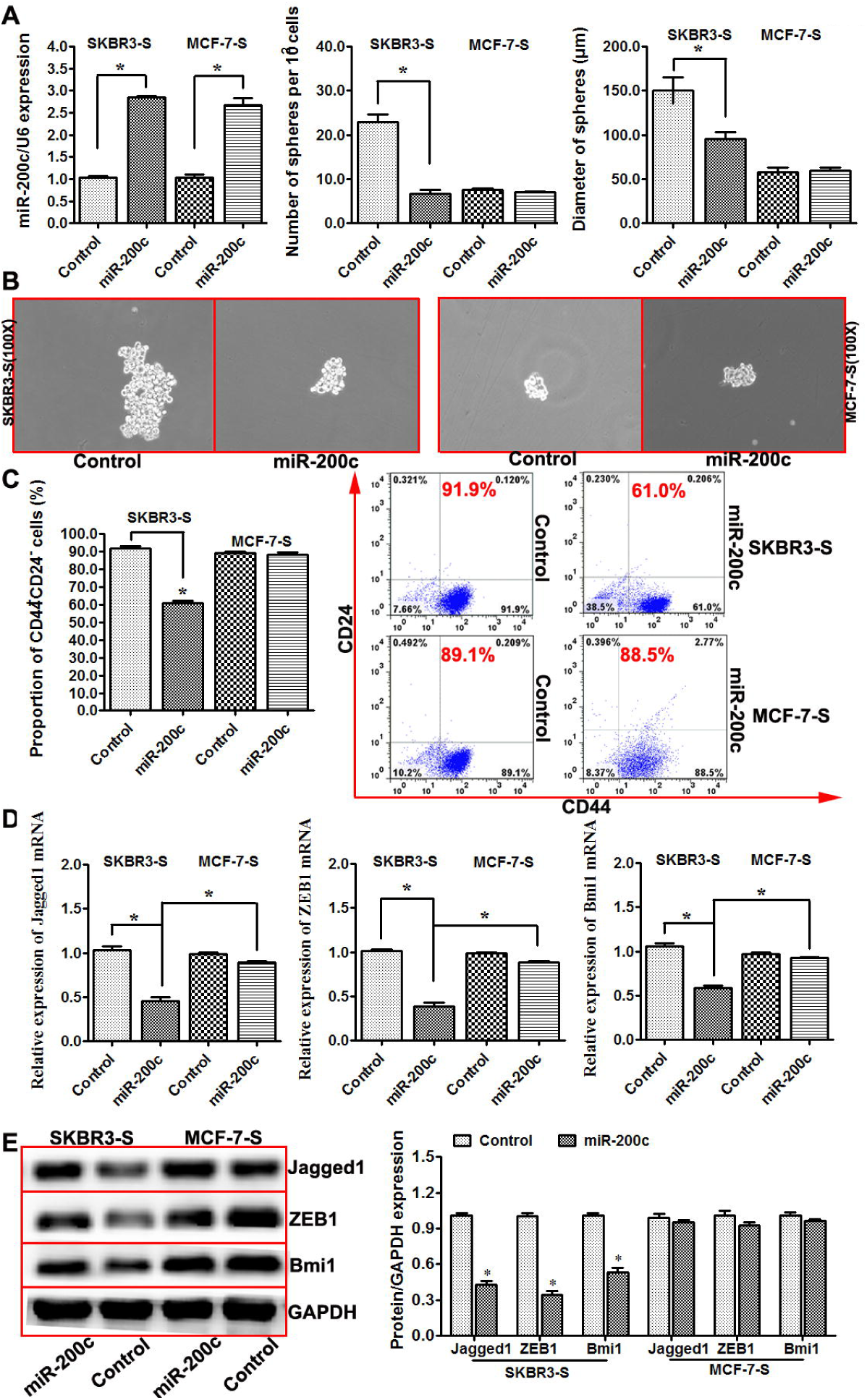
Overexpression of miR-200c resulted in a decrease in the number of tumor spheres and proportion of CD44^+^CD24^-^ phenotype cells in HER2+ BCSCs. (A) The number and diameter of tumor spheres formed per 100 cells from SKBR3-S and MCF-7-S. First, SKBR3-S and MCF-7-S were stably transfected with lentiviral vector encoding miR-200c or its negative control. The transfection was successful. The spheres were counted for 10 fields in each group. miR-200c overexpression resulted in an obvious decrease in the number and diameter of tumor spheres formed from SKBR3-S but not MCF-7-S (n=3, *p<0.05). (B) Representative images showing tumor spheres formed from SKBR3-S and MCF-7-S cells expressing the indicated constructs (magnification of 100x). (C) Flow cytometric profiles of CD44^+^CD24^-^ phenotype cells among SKBR3-S and MCF-7-S cells after stable expression of miR-200c. Overexpression of miR-200c reduced the proportion of CD44^+^CD24^-^ phenotype cells from SKBR3-S but not MCF-7-S (n=3, *p< 0.05). (D) qRT-PCR was used to detect the expression of Jagged1, ZEB1 and Bmi1 mRNAs in SKBR3-S and MCF-7-S cells after stable expression of miR-200c. The expression levels of Jagged1, ZEB1 and Bmi1 mRNAs were decreased when transfected with lentiviral vector encoding miR-200c in SKBR3-S but not MCF-7-S (*p<0.05 vs. ControlSKBR3-S), and their expression levels were significantly lower in SKBR3 than that in MCF-7 after stable expression of miR-200c (*p<0.05 vs. miR-200cMCF-7). (E) Western blot was used to detect the protein expression of Jagged1, ZEB1 and Bmi1 in SKBR3-S and MCF-7-S cells after stable expression of miR-200c. The expression of Jagged1, ZEB1 and Bmi1 proteins were decreased when transfected with lentiviral vector encoding miR-200c in SKBR3-S but not MCF-7-S (*p<0.05 vs. Control).

Our results demonstrated that overexpression of miR-200c resulted in an obvious decrease in the number and diameter of tumor spheres formed from sorted CD44^+^CD24^-^ phenotype SKBR3-S but not MCF-7-S cells (Figure 2A). Images of tumor spheres formed from SKBR3-S and MCF-7-S cells transfected with lentiviral vector are shown (100X, Figure 2B). Flow cytometry was used to detect the proportion of CD44^+^CD24^-^ phenotype cells, and the results showed that overexpression of miR-200c reduced the proportion of CD44^+^CD24^-^ phenotype SKBR3-S cells but not MCF-7-S cells (Figure 2C). Then, we examined whether overexpression of miR-200c modulates Jagged1, ZEB1 and Bmi1 expression in HER2+ stem cells. qRT-PCR and western blot results showed that the expression levels of Jagged1, ZEB1 and Bmi1 mRNAs and proteins were decreased when transfected with lentiviral vector encoding miR-200c in SKBR3-S but not MCF-7-S cells (Figure 2D, 2E).

### miR-200c reduced the stemness and increased the sensitivity to trastuzumab in HER2+ breast cancer cells and stem cells

First, SKBR3-S and MCF-7-S cells were treated with 0 μg/ml or 10 μg/ml trastuzumab. Then, we assessed the number and diameter of tumor spheres formed per 100 cells from SKBR3-S and MCF-7-S cells. The number and diameter were noticeably decreased in SKBR3-S cells but not MCF-7-S cells after treatment with 10 μg/ml trastuzumab (Figure 3A). Next, we assessed the proportion of CD44^+^CD24^-^ phenotype cells and found that the proportion was also noticeably decreased in SKBR3-S but not MCF-7-S cells after treatment with 10 μg/ml trastuzumab (Figure 3B). Images of tumor spheres formed from SKBR3-S and MCF-7-S cells treated with or without trastuzumab are shown (100X, Figure 3C). Then, we detected the expression of miR-200c in SKBR3-S, SKBR3, MCF-7-S and MCF-7 cells with or without the treatment of 10 μg/ml trastuzumab. We found that expression of miR-200c was significantly increased in SKBR3-S and SKBR3 but not in MCF-7-S and MCF-7 (Figure 3D), indicating that upon treatment with trastuzumab, the expression of miR-200c was up-regulated only in HER2+, not HER2-, breast cancer cells and stem cells.

**Figure 3.**
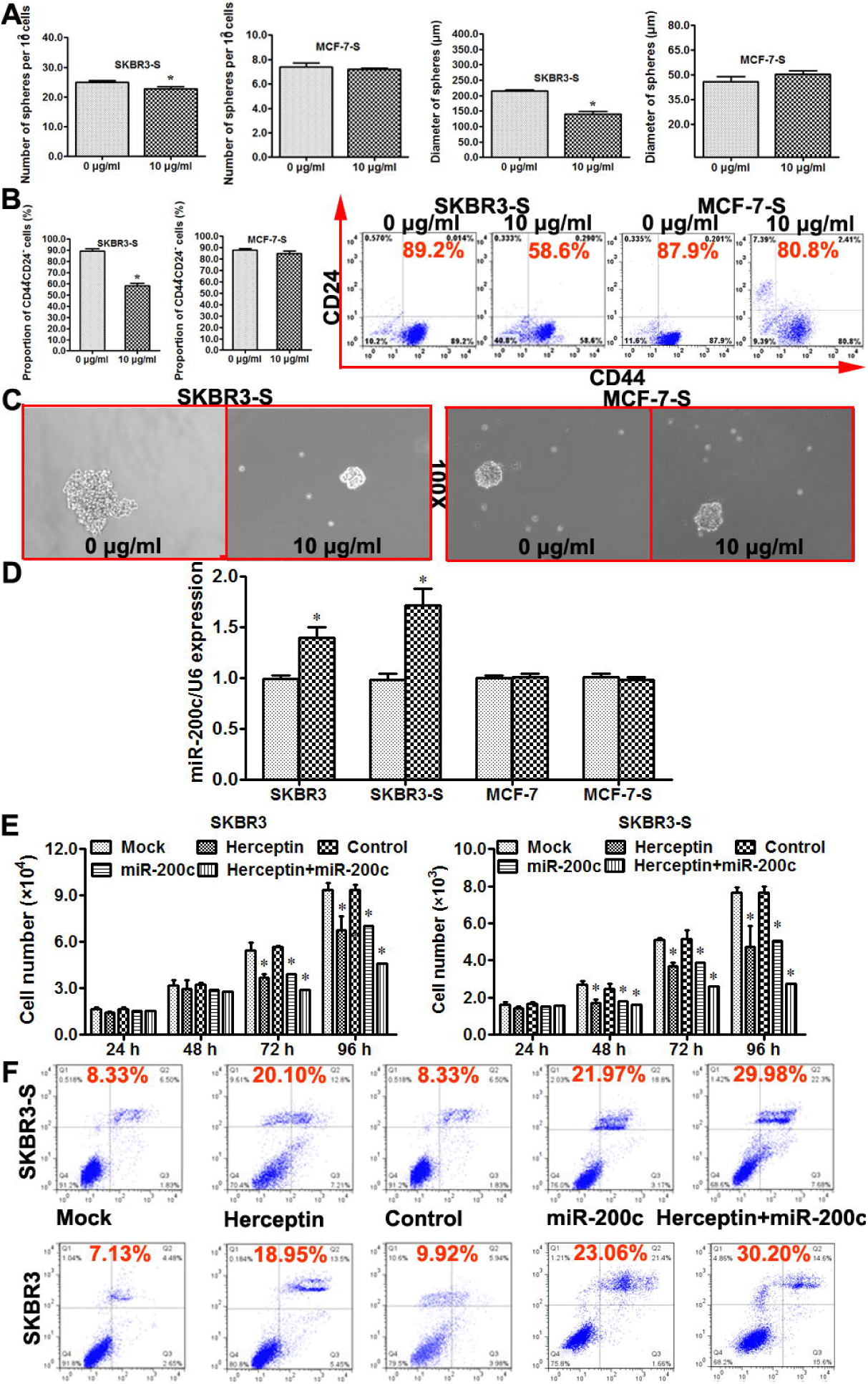
miR-200c reduced the stemness and increased the sensitivity to trastuzumab in HER2+ breast cancer cells and stem cells. (A) The number and diameter of tumor spheres formed per 100 cells from SKBR3-S and MCF-7-S cells. SKBR3-S and MCF-7-S cells were treated with trastuzumab at 0 μg/ml or 10 μg/ml. Trastuzumab was administered for 3 days and refreshed trastuzumab every 24 h. The spheres were counted for 10 fields in each group. Trastuzumab treatment increased the number and diameter of tumor spheres formed per 100 cells from SKBR3-S but not MCF-7-S 72 hours after trastuzumab treatment (n=3, *p< 0.05). (B) Flow cytometric profiles of CD44^+^CD24^-^ phenotype cells for SKBR3-S and MCF-7-S cells after 0 μg/ml or 10 μg/ml trastuzumab treatment. The proportion was noticeably decreased in the 10 μg/ml trastuzumab treatment group of SKBR3-S but not MCF-7-S cells (n=3, *p< 0.05). (C) Representative images showing tumor spheres formed from SKBR3-S and MCF-7-S cells with 0 μg/ml or 10 μg/ml trastuzumab treatment (magnification of 100x). (D) qRT-PCR was used to evaluated the expression of miR-200c in SKBR3, SKBR3-S, MCF-7 and MCF-7-S after 0 μg/ml or 10 μg/ml trastuzumab treatment (n=3, *p< 0.05). (E) MTT assay was used to assess the proliferation of SKBR3 and SKBR3-S in different groups. SKBR3 and SKBR3-S cells were divided into five groups: Mock (blank control), Trastuzumab (10 μg/ml), Control (transiently transfected with lentiviral vector encoding the negative control of miR-200c), miR-200c (transiently transfected with lentiviral vector encoding miR-200c), and Trastuzumab+miR-200c. The proliferation of SKBR3 and SKBR3-S was significantly decreased in the Trastuzumab, miR-200c and Trastuzumab+miR-200c groups (*p<0.05 vs. Mock, *p<0.05 vs. Control), with the largest decrease in the Trastuzumab+miR-200c group (*p<0.05 vs. miR-200c, *p<0.05 vs. Trastuzumab). (F) PI/Annexin V double staining with FCM was used to detect the percentage of apoptotic cells in SKBR3 and SKBR3-S cells. SKBR3 and SKBR3-S cells were divided into five groups as for the MTT assay: Mock, Trastuzumab, Control, miR-200c, and Trastuzumab+miR-200c. The percentage of apoptotic cells was significantly increased in the Trastuzumab, miR-200c and Trastuzumab+miR-200c groups (*p<0.05 vs. Mock, *p<0.05 vs. Control), with the largest increase in the Trastuzumab+miR-200c group (*p<0.05 vs. miR-200c, *p<0.05 vs. Trastuzumab).

Subsequently, we divided SKBR3 and SKBR3-S cells into mock (negative control), Trastuzumab (10 μg/ml), Control (transiently transfected with lentiviral vector coding control), miR-200c (transiently transfected with lentiviral vector coding miR-200c), and Trastuzumab+miR-200c groups. The MTT assay and PI/Annexin V double staining with the FCM assay were performed to detect the proliferation rate of these cells and the percentage of apoptotic cells in SKBR3 and SKBR3-S. The proliferation ability of SKBR3 and SKBR3-S was significantly decreased in the Trastuzumab, miR-200c and Trastuzumab+miR-200c groups, with the largest decrease in the Trastuzumab+miR-200c group (Figure 3E). Furthermore, the percentage of apoptotic cells in SKBR3 and SKBR3-S was significantly increased in the Trastuzumab, miR-200c and Trastuzumab+miR-200c groups, with the largest increase in the Trastuzumab+miR-200c group (Figure 3F).

### miR-200c significantly inhibited tumor growth and metastasis of HER2+ breast cancer cells and stem cells in vivo

To explore the role of miR-200c for BCSC function in vivo, we performed xenograft experiments with SKBR3, SKBR3-S, MCF-7 and MCF-7-S cells that were stably transfected with lentiviral vector coding miR-200c or its negative control. These cells were injected subcutaneously into the flanks of Balb/c mice (five in each group). The tumor volume was measured and recorded every four days after injection (tumor volume=long diameter*short axis square/2). After 28 days, all mice were sacrificed to harvest tumors. The growth curves of the tumor volume were plotted. The final weights of tumors were recorded. Images of subcutaneous tumors from SKBR3, SKBR3-S, MCF-7 and MCF-7-S cells expressing the indicated constructs are shown in Figure 4A. The results demonstrated that miR-200c significantly reduces the volume and weight of tumors from SKBR3-S and SKBR3 cells, whereas there was almost no decrease in the volume and weight of tumors from MCF-7-S and MCF-7 (Figure 4B, 4C). The mice were also dissected for lung tissues to observe the number of tumor metastases. Lung tissues and their HE staining images from SKBR3, SKBR3-S, MCF-7 and MCF-7-S are shown in the left panel of Figure 4D. We found that miR-200c prevents metastasis in SKBR3-S and SKBR3 cells. The number of lung metastases decreased significantly after stable transfection with lentiviral vector coding miR-200c in SKBR3-S and SKBR3 cells, whereas no such changes were observed in MCF-7-S and MCF-7 cells (Figure 4D).

**Figure 4.**
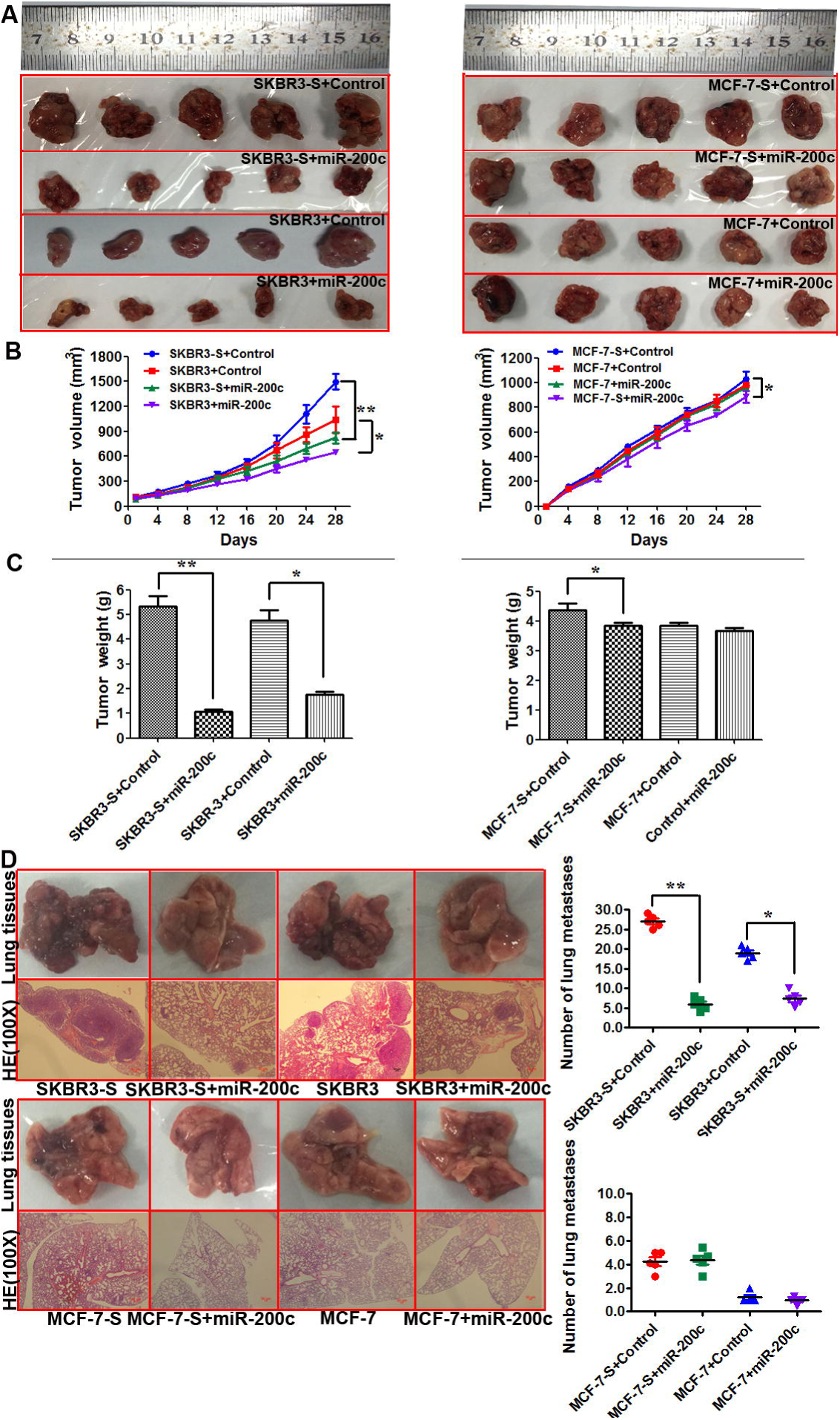
miR-200c significantly inhibited tumor growth and metastasis of HER2+ breast cancer cells and stem cells in vivo. (A) Images of subcutaneous tumors from SKBR3, SKBR3-S, MCF-7 and MCF-7-S cells that were stably transfected with lentiviral vector coding miR-200c or control. (B) Tumor volume in mouse xenograft models. SKBR3, SKBR3-S, MCF-7 and MCF-7-S cells expressing the indicated constructs were injected subcutaneously into BALC/c mice (five in each group). The tumor volume was measured every four days. miR-200c significantly reduced the volume of tumors from SKBR3 and SKBR3-S cells (*p<0.05 vs. Control), whereas there was almost no decrease in the volume of tumors from MCF-7-S and MCF-7 cells. (C) Tumor weight in mouse xenograft models. After 28 days, the mice were killed, necropsies were performed, and the tumors were weighed. miR-200c significantly reduced the weight of tumors from SKBR3 and SKBR3-S cells (*p<0.05 vs. Control), whereas there was almost no decrease in the weight of tumors from MCF-7-S and MCF-7 cells. (D) Lung tissues and their HE staining images from SKBR3, SKBR3-S, MCF-7 and MCF-7-S cells that were stably transfected with lentiviral vector coding miR-200c or control (*p<0.05 vs. Control).

### Jagged1, ZEB1 and Bmi1 were highly expressed and positively related with each other in clinical specimens from breast cancer patients

All tissue samples in this study were collected from 50 breast cancer patients. The breast cancer tissues (Breast cancer) and their matched adjacent tissues (Normal) were removed during surgery and immediately stored in RNA later for RNA extraction and subsequent qRT-PCR assay. qRT-PCR results revealed that Jagged1, ZEB1 and Bmi1 mRNAs were upregulated in breast cancer tissues compared with matched adjacent tissues (Figure 5A).

**Figure 5.**
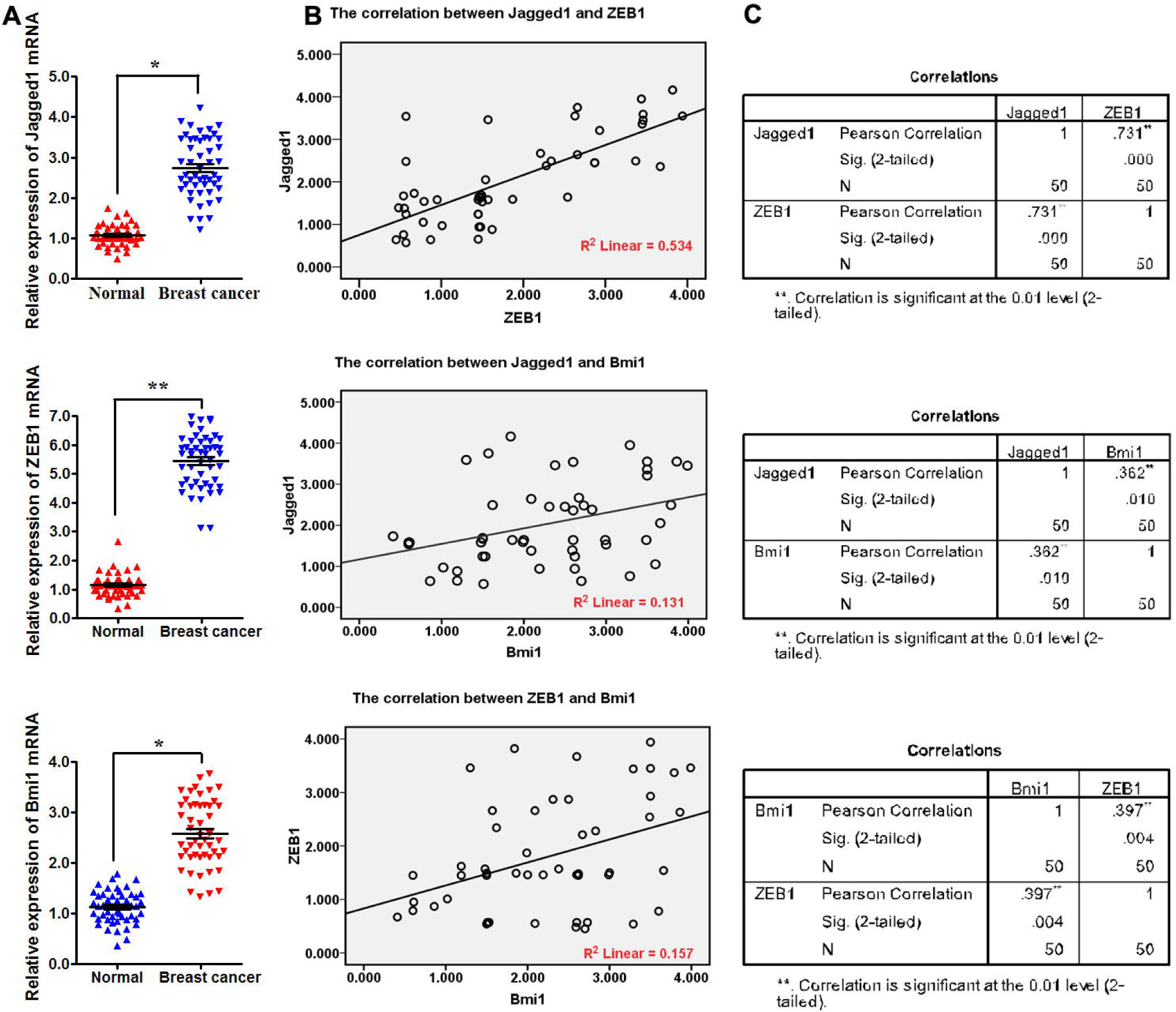
Jagged1, ZEB1 and Bmi1 were highly expressed and positively related with each other in clinical specimens from breast cancer patients. (A) qRT-PCR was used to detect the expression of Jagged1, ZEB1 and Bmi1 mRNAs in breast cancer patients (*p<0.05 vs. Normal). (B) Simple scatter plot of the Pearson correction analysis between the expression of Jagged1, ZEB1 and Bmi1 mRNAs in breast cancer patients. (C) The expression levels of Jagged1 and ZEB1 were more strongly related than those of Jagged1 and Bmi1 or ZEB1 and Bmi1 (2-tailed).

Then, we explored the correlation between Jagged1, ZEB1 and Bmi1 expression using Pearson correlation analysis. The correlation coefficient between Jagged1 and ZEB1 was 0.731 (linear R2 = 0.534), between Jagged1 and Bmi1 was 0.362 (linear R2 = 0.131), and between ZEB1 and Bmi1 was 0.397 (linear R2 = 0.157), indicating that the expression levels of Jagged1 and ZEB1 were more strongly related than those of Jagged1 and Bmi1 or ZEB1 and Bmi1 (2-tailed, Figure 5B, 5C).

### Mechanisms of trastuzumab resistance in HER2-overexpressing breast cancer

miR-200c inhibits stemness and increases the sensitivity of HER2+ cells and stem cells to trastuzumab via the Notch, Wnt and Hedgehog pathways. miR-200c inhibits the activity of the Notch pathway by Jagged1, which is the ligand of the Notch pathway leading to HER2 overexpression and stemness enhancement of BCSCs. miR-200c inhibits the Wnt pathway by ZEB1, which can lead to metastasis and resistance in breast cancer. miR-200c inhibits the Hedgehog pathway by Bmi1, which is a known regulator of stem cell self-renewal and tumor growth. Jagged1, ZEB1 and Bmi1 are regulated by miR-200c. By directly targeting these three genes, miR-200c coordinates the stemness-related Notch, Wnt and Hedgehog signaling pathways. Finally, miR-200c affects stem cell properties and drug resistance in HER2-overexpressing breast cancer (Figure 6).

**Figure 6.**
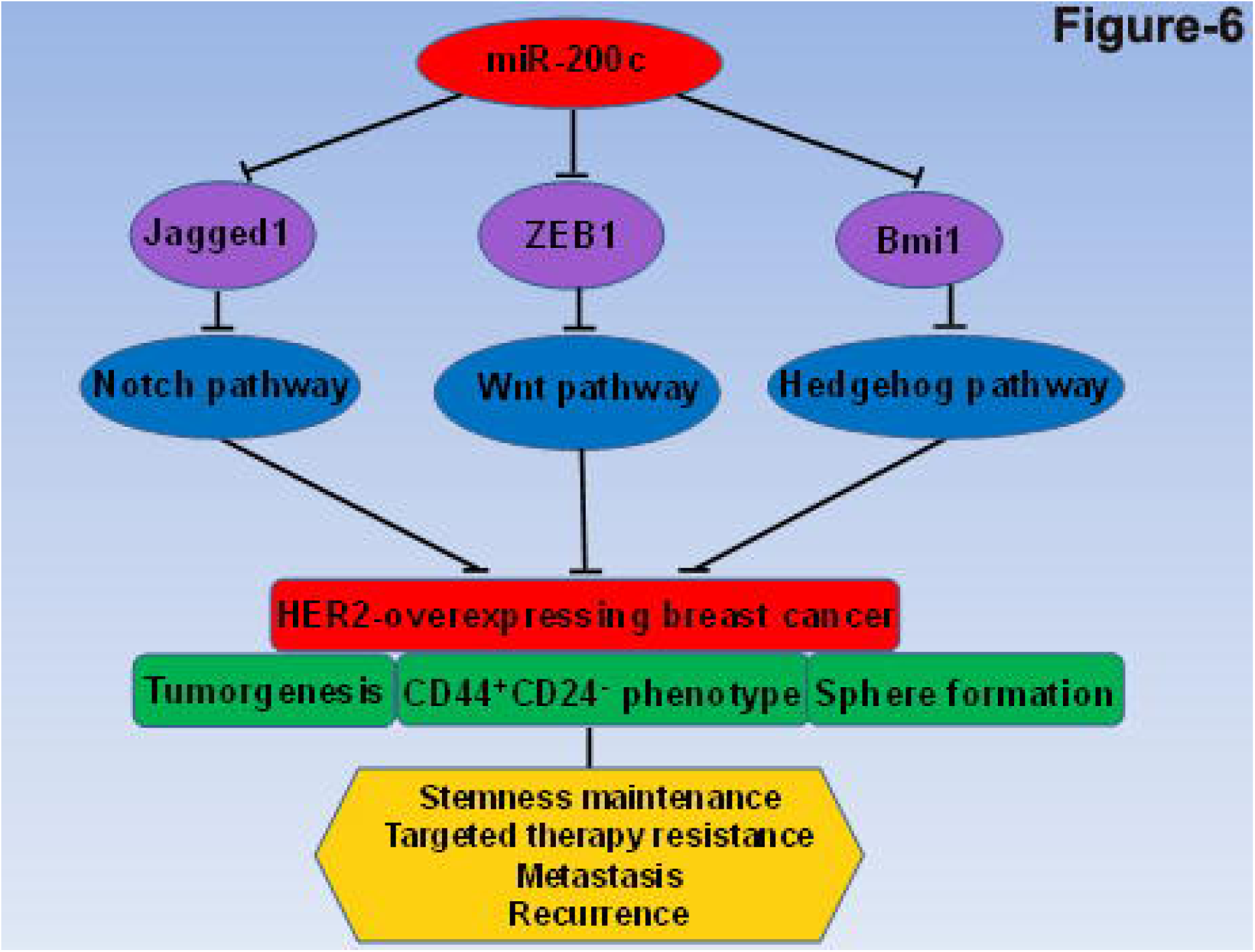
Mechanisms of trastuzumab resistance in HER2-overexpressing breast cancer. By directly targeting Jagged1, ZEB1 and Bmi1, miR-200c coordinates the stemness-related Notch, Wnt and Hedgehog signaling pathways. Finally, miR-200c affects stem cell properties and drug resistance in HER2-overexpressing breast cancer.

## Discussion

Trastuzumab has significantly improved the overall survival of HER2-overexpressing breast cancer patients. However, more than 70% of trastuzumab-treated patients have experienced disease progression or relapse in past years. Inherent and acquired resistance to chemotherapy, radiotherapy or targeted therapy among the CSC population can be used to explain why conventional therapies fail and tumor relapses occur. Because of insufficient targeting to BCSCs, trastuzumab treatment finally fails (Nahta and Esteva, 2006; Valabrega et al., 2007). Therefore, treatment for HER2-overexpressing breast cancer should target the BCSC population along with the remaining tumor cells. Many studies suggested that the Notch, Wnt and Hedgehog pathways play an extremely important role in stemness maintenance of BCSCs and treatment resistance (Li et al., 2003; Merchant and Matsui, 2010; Pannuti et al., 2010). At the same time, miR-200c is often weakly expressed in cancer and is closely related to these signal pathways by directly targeting three important genes: Jagged1, ZEB1 and Bmi1 (Korpal et al., 2008; Liu et al., 2014; Vallejo et al., 2011). We hypothesized that miR-200c can suppress stemness and increase the cellular sensitivity to trastuzumab in HER2+ breast cancer cells and stem cells via Jagged1, ZEB1 and Bmi1.

The biological role of miR-200c in tumor cells and stem cells has been demonstrated in many cancers (Shimono et al., 2009). To further explore the role of miR-200c in HER2-overexpressing breast cancer, we first compared the expression of miR-200c in 9 breast cancer cells and 1 mammary epithelial cell line and in CD44^+^CD24^-^ phenotype stem cells of the above 10 cell lines. We found that miR-200c was weakly expressed in both breast cancer cells and CD44^+^CD24^-^ phenotype stem cells. Then, we detected the expression of miR-200c in SKBR3-S/SKBR3-nS, MCF-7-S/MCF-7-nS, SKBR3-SM/SKBR3-nSM, MCF-7-SM/MCF-7-nSM and normal adherent cells of SKBR3 and MCF-7. The results showed that miR-200c was more weakly expressed in SKBR3-S and SKBR3-SM than SKBR3 and was the lowest in SKBR3-S. However, there were no significant differences between the expression of miR-200c in MCF-7, MCF-7-S and MCF-7-SM, suggesting that miR-200c may have a greater functional impact on purified HER2+ SKBR3-S than HER2-MCF-7-S. Down-regulation of miR-200c has been reported in many tumors and is related to the occurrence and development of different tumors (Iliopoulos et al., 2010). Here, we first report that down-regulation of miR-200c is much greater in the CD44^+^CD24^-^ phenotype BCSCs and suspension microspheres than in normal adherent cells of HER2+ breast cancer cells and that the most significant down-regulation occurs in HER2+ CD44^+^CD24^-^ phenotype stem cells. Based on these results, we hypothesize that down-regulation of miR-200c in HER2+ breast cancer cells is correlated with stemness maintenance.

Then, we stably overexpressed miR-200c in SKBR3-S and MCF-7-S to observe stemness-related changes. We found that the number and diameter of tumor spheres formed per 100 cells and the proportion of CD44^+^CD24^-^ phenotype cells were significantly reduced in SKBR3-S but not MCF-7-S cells after stable transfection of a lentiviral vector encoding miR-200c. These results demonstrate that miR-200c has a greater inhibitory effect on stem cells from HER2+ breast cancer cells than HER2-cells. We further validated this conclusion in HER2+ and HER2-stem cells by western blot and qRT-PCR. The mRNA and protein expression levels of Jagged1, ZEB1 and Bmi1 were significantly decreased. Studies showed that Jagged1, ZEB1 and Bmi1 are three important genes, respectively, in the Notch, Wnt and Hedgehog pathways that induce pluripotent stem cells. They all have a binding site for miR-200c at the 3’-UTR end. The expression of Jagged1, ZEB1 and Bmi1 is correlated with tumor stem cells (Korpal et al., 2008; Liu et al., 2014; Vallejo et al., 2011). Therefore, the decrease in mRNA and protein expression of Jagged1, ZEB1 and Bmi1 after overexpression of miR-200c was consistent with these reports.

Then, we treated SKBR3-S and MCF-7-S cells with trastuzumab at 0 μg/ml or 10 μg/ml. The results showed that the number and diameter of tumor spheres formed and the proportion of CD44^+^CD24^-^ phenotype cells from SKBR3-S were noticeably decreased. The expression of miR-200c was noticeably increased in the 10 μg/ml trastuzumab-treated group of SKBR3-S, and SKBR3 but not MCF-7-S and MCF-7 cells. This indicates that upon treatment with trastuzumab, the stemness of HER2+ BCSCs decreased and the expression of miR-200c increased. This may be the key to inhibiting tumor recurrence and resistance in HER2+ breast cancer. The MTT assay showed that the proliferation of SKBR3 and SKBR3-S was significantly decreased in the Trastuzumab, miR-200c and Trastuzumab+miR-200c groups, with the largest decrease in the Trastuzumab+miR-200c group. PI/Annexin V double staining with the FCM assay showed that the percentage of apoptotic cells in SKBR3 and SKBR3-S was significantly increased in the Trastuzumab, miR-200c and Trastuzumab+miR-200c groups, with the largest increase in the Trastuzumab+miR-200c group. These results demonstrate that combining miR-200c with trastuzumab can significantly reduce proliferation and increase apoptosis of HER2+ breast cancer cells and stem cells. These results first verify that miR-200c plays a vital role in stemness maintenance and increases the sensitivity of HER2+ breast cancer cells and stem cells to trastuzumab.

Then, we found that miR-200c reduced HER2+ breast cancer cells and stem cell progression into malignant tumors in xenograft experiments. These results were consistent with the in vitro cell experiments. miR-200c blocked tumor formation in nude mice from HER2+ cells and stem cells, including reducing the tumor volume, size, weight and lung metastasis. These results are consistent with reports that miR-200c overexpression improves the overall survival of tumor patients (Leskela et al., 2011; Song et al., 2015). Finally, we found that Jagged1, ZEB1 and Bmi1 were highly expressed and positively related with each other in clinical specimens from breast cancer patients. As has been confirmed, miR-200c coordinates several stemness-related signaling cascades, such as Notch, Wnt and Hedgehog. Low levels of miR-200c affect stem cell properties and drug resistance in breast cancer (Liu and Tang, 2011). The activated Notch signaling pathway plays a vital role in controlling the stem-like phenotype of stem cells. Notch receptors and their ligands are aberrantly activated in many types of human cancers. miR-200c can activate the Notch signaling pathway by targeting the ligand of Notch (Jagged-1) (Gallardo et al., 2015). Additionally, miR-200c is associated with the increased expression of ZEB-1, which is the direct target gene of miR-200c, and important genes in stemness-related Wnt pathways (Ramachandran et al., 2014). Bmi1 is a key member of the Hedgehog pathway. Bmi1 is a regulator of stem cell self-renewal and suppresses the mammary duct formation ability of mammary stem cells and tumor growth driven by BCSCs (Molofsky et al., 2003). Jagged1, ZEB1 and Bmi1 are regulated by miR-200c. The expression of Jagged1, ZEB1 and Bmi1 increased after the abnormally low expression of miR-200c. These findings show the connection between miR-200c, Notch, Wnt and Hedgehog signaling in HER2 overexpressing breast cancer.

In conclusion, these results are consistent with our hypothesis that miR-200c reduces stemness and increases sensitivity to trastuzumab in HER2+ breast cancer cells and stem cells via inhibition of Jagged1, ZEB1 and Bmi1, which are three important genes in the stemness-related Notch, Wnt and Hedgehog pathways. However, many molecular mechanisms remain unclear. We will continue to further explore the detailed molecular mechanisms and biological and pathological functions of miR-200c in increasing drug sensitivity to trastuzumab via these three pathways.

## Acknowledgements

We are grateful to all the SYSUCC colleagues, especially Dr. Bo Chen and Xiaojia Huang, for their permanent support, inspiration, constructive discussions and critical reading of the manuscript.

## Authors’ Contributions

C. Song, H. Tang and X, Xie designed the experiments. C. Song interpreted the data and wrote the manuscript. C. Song and H. Tang carried out experiments. J. Wang, H. Wang, P. Liu, L. Liu and L. Yang collected the human samples and clinical data.

## Funding

This work was supported by funds from the National Natural Science Foundation of China (81472469, to Hailin Tang; 81472575 and 81672598, to Xiaoming Xie).

The costs of publication of this article were defrayed in part by the payment of page charges. This article must therefore be hereby marked advertisement in accordance with 18 U.S.C. Section 1734 solely to indicate this fact.

